# Interaction between Solo and PDZ-RhoGEF is involved in actin cytoskeletal remodeling and response to substrate stiffness

**DOI:** 10.1101/2023.11.08.566199

**Authors:** Aoi Kunitomi, Shuhei Chiba, Nahoko Higashitani, Atsushi Higashitani, Shinichi Sato, Kensaku Mizuno, Kazumasa Ohashi

## Abstract

Recent findings indicate that Solo, a RhoGEF, is involved in cellular mechanical stress responses. However, the mechanism of actin cytoskeletal remodeling via Solo remains unclear. Therefore, this study was aimed at identifying Solo-interacting proteins using the BioID, a proximal-dependent labeling method and elucidating the molecular mechanisms of function of Solo. We identified PDZ-RhoGEF (PRG) as a Solo-interacting protein. PRG co-localized with Solo in the basal area of cells, depending on Solo localization, and enhanced actin polymerization at Solo accumulation sites. Additionally, Solo and PRG interaction was necessary for actin cytoskeletal remodeling and RhoA activation. Moreover, overexpression of the binding domains of Solo and PRG had a dominant-negative effect on actin polymerization and actin stress fiber formation in response to substrate stiffness. Therefore, Solo restricts the localization of PRG and regulates actin cytoskeletal remodeling in synergy with PRG in response to the surrounding mechanical environment.

## INTRODUCTION

The regulation of actin cytoskeletal remodeling has diversified during the evolution of multicellular organisms. In the process, many actin regulatory proteins and signaling pathways are activated to generate highly spatiotemporally regulated actin structures (dos Remedios *et al*., 2003; Blanchoin *et al*., 2014; Pollard and Goldman, 2018). Rho family small GTPases, encoded by approximately 20 genes in the human genome, are key regulators of actin cytoskeletal remodeling (Jaffe and Hall, 2005; Heasman and Ridley, 2008). Members of the Rho family have a GDP-bound inactive form and a GTP-bound active form. Rho GTPases must be tightly regulated by upstream signaling molecules to construct the appropriate high-order actin structures in response to various stimuli (Bos *et al*., 2007). Approximately 80 Rho guanine nucleotide exchange factors (RhoGEFs, activators of Rho family small GTPases) and RhoGAPs (their inactivators) are encoded by the human genome. Molecular diversity enables the spatiotemporal regulation of Rho GTPases in response to various external stimuli (Tcherkezian and Lamarche-Vane, 2007; Cherfils and Zeghouf, 2013; Cook *et al*., 2014). Recent findings suggest a crosstalk between the upstream regulators of Rho GTPases, including RhoGEF and RhoGAP, and this crosstalk is thought to precisely regulate Rho GTPases (Guilluy *et al*., 2011a; Hodge and Ridley, 2016).

Actin cytoskeletal remodeling is important for mechanical stress response in cells involved in embryogenesis, tissue morphogenesis, and various physiological phenomena (Lecuit *et al*., 2011; Romet-Lemonne and Jegou, 2013). Notably, some RhoGEFs are involved in mechanical stress response. For example, leukemia-associated RhoGEF (LARG, ARHGEF12) and GEF-H1 (ARHGEF2) are required for tensile force-induced RhoA activation in cells via integrins (Guilluy *et al*., 2011b). p114RhoGEF (ARHGEF18) and PDZ-RhoGEF (ARHGEF11, hereafter designated as PRG) accumulate at cell-cell adhesions in a force-dependent manner and regulate actin cytoskeletal remodeling through RhoA activation (Ito *et al*., 2017; Acharya *et al*., 2018). Moreover, 11 RhoGEFs, including LARG and GEF-H1, are involved in cyclic stretching-induced cell alignment (Abiko *et al*., 2015). Notably, Solo (ARHGEF40), one of the 11 RhoGEFs, is required for tensile force-induced RhoA activation and reinforcement of stress fibers in cells (Fujiwara *et al*., 2016). Therefore, Solo is believed to play a crucial role in mechanical stress response in cells. However, the molecular mechanisms that regulate the localization and activity of Solo remain unknown.

In this study, we comprehensively analyzed for Solo interacting proteins using proximity-dependent biotin identification (BioID) coupled with proteomics (Roux *et al*., 2012). We provide evidence that PRG binds to Solo and the relationship between Solo and PRG cooperatively regulates actin cytoskeletal remodeling and is involved in the response of cells to substrate stiffness.

## RESULTS

### Identification of PDZ-RhoGEF as a Solo-interacting protein

To elucidate the mechanisms underlying the localization and function of Solo, we searched for Solo-interacting proteins using the TurboID-based proximity-dependent biotin-labeling method (Branon et al., 2018). TurboID is an engineered form of biotin ligase (BirA), which can label the proteins proximal to the TurboID-fused protein more efficiently and rapidly than BirA*. BirA* is a mutated BirA that loses substrate specificity. (Roux *et al*., 2012). We constructed expression plasmids coding for Solo N-or C-terminal fused with the V5 epitope and TurboID (Figure 1A; V5-TurboID-Solo and Solo-TurboID-V5) and established human colorectal adenocarcinoma cell lines (DLD1) stably expressing V5-and TurboID-fused Solo or control V5-TurboID. These cell lines and their parental cells were treated with biotin for 4 h and lysed, and the biotin-labeled proteins in the cell lysates were precipitated with tamavidin-conjugated beads. Tamavidin is an avidin-like, biotin-binding protein derived from mushrooms (Takakura *et al*., 2009). The precipitated proteins were separated using sodium dodecyl sulfate-polyacrylamide gel electrophoresis (SDS-PAGE) and stained with silver (Figure 1B). Individual bands were excised, trypsinized, and analyzed using mass spectrometry. Actin and several actin-related proteins were identified as candidate proteins localized in proximity to Solo (Figure 1C), including two RhoGEFs (PDZ-RhoGEF [also known as ARHGEF11] and LARG [also known as ARHGEF12]), unconventional myosins (myosin VI and members of the myosin I family), and spectrin. Although studies have confirmed that PRG and LARG are Solo-binding proteins, the role of their interactions in cells remains to be explored (O’Loughlin *et al*., 2018; Muller *et al*., 2020). Intriguingly, three RhoGEFs–Solo, PRG, and LARG are involved in mechanical stress responses (Guilluy *et al*., 2011b; Fujiwara *et al*., 2016; Ito *et al*., 2017). Therefore, co-immunoprecipitation assay was performed to examine the interaction between Solo and PRG. COS-7 cells were transfected with yellow fluorescent protein (YFP)-tagged Solo, followed by immunoprecipitation of the cell lysates with an anti-GFP nanobody and immunoblotting of the precipitates with anti-GFP and anti-PRG antibodies. Endogenous PRG co-precipitated with YFP-Solo, but not with control YFP (Figure 1D), indicating that PRG specifically interacted with Solo in the cells.

**FIGURE 1:**
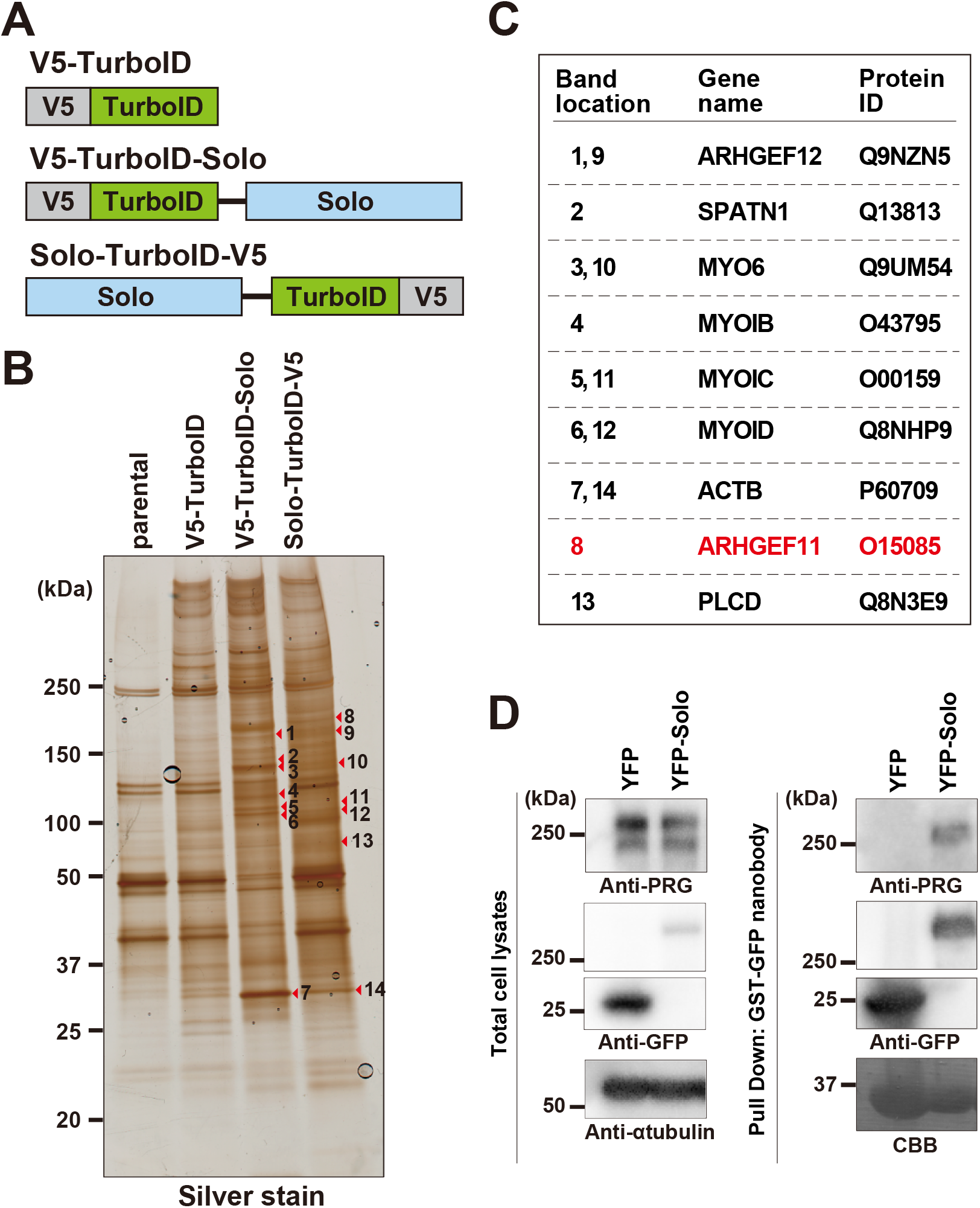
Identification of candidates for Solo-interacting proteins using TurboID and mass spectrometry. (A) Schematic structures of V5-TurboID, V5-TurboID-Solo, and Solo-TurboID-V5. (B) DLD-1 cells stably expressing the TurboID and TurboID-fused proteins were cultured in a biotin-containing culture medium and then lysed. Cell lysates were precipitated with streptavidin beads. The precipitated biotinylated proteins were separated using SDS-PAGE, detected via silver staining, and identified using mass spectrometric analysis. (C) A partial list of identified proteins. The numbers on the list correspond to the numbers on the silver-stained gel in B. (D) Co-immunoprecipitation assay of Solo and PRG. COS-7 cells were transfected with YFP or YFP-Solo. Cell lysates were immunoprecipitated with anti-GFP nanobody, and precipitated proteins were analyzed by immunoblotting with anti-GFP, anti-PRG and anti-α-tubulin antibodies. GST-nanobody was detected with CBB.

### PRG co-localizes with Solo and promotes Solo-induced actin assembly

To examine the subcellular localization of Solo and PRG, YFP-tagged Solo and mCherry-tagged PRG were individually expressed in Madin-Darby canine kidney (MDCK) cells, and their localization was analyzed using fluorescence microscopy (Figure 2A). Actin filaments were visualized by staining with Alexa Fluor 647-conjugated phalloidin (Alexa 647-phalloidin). Consistent with our previous findings (Fujiwara *et al*., 2018), confocal microscopy of the basal plane of the cells revealed that YFP-Solo was uniquely localized to punctate dots, which were often aligned in a direction perpendicular to stress fibers on the basal plane of the cells. Additionally, YFP-Solo expression moderately promoted stress fiber formation in the whole cell and local accumulation of actin filaments at Solo accumulation sites (Figure 2A). On the other hand, mCherry-PRG was localized to both the cell periphery and punctate dots on the basal plane, and its expression promoted lamellipodium-like actin assembly at the cell periphery and stress fiber formation on the basal plane in accordance with previous results showing that PRG has the potential to activate RhoA and Cdc42 (Figure 2A, Banerjee *et al*., 2009; Castillo-Kauil *et al*., 2020). When YFP-Solo and mCherry-PRG were co-expressed in MDCK cells, PRG co-localized with Solo at the unique linear alignment on the basal plane, with a decrease in the localization of PRG at lamellipodia (Figure 2A). This result suggests that Solo guides PRG to the site of Solo localization on the basal plane of the cells.

**FIGURE 2:**
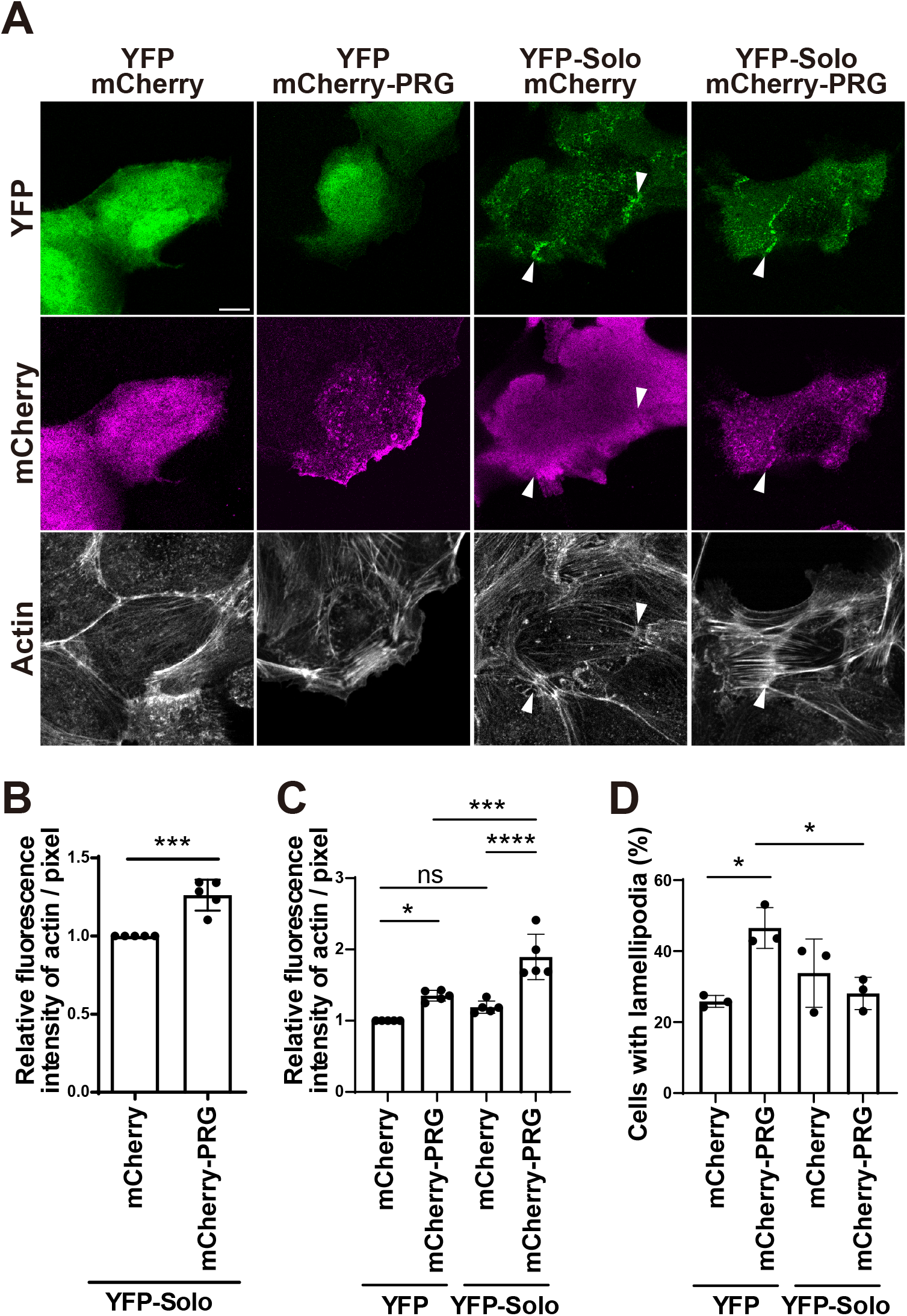
PRG co-localizes with Solo and enhances Solo-induced actin polymerization. (A) Confocal microscopic images of YFP or YFP-Solo, mCherry or mCherry-PRG, and F-actin in MDCK cells. MDCK cells transfected with the indicated plasmids and CFP-CAAX expression plasmid were fixed and stained with anti-mCherry (magenta) and phalloidin (gray). White arrowheads indicate the accumulation sites of YFP-Solo in the basal area. Scale bar, 10 µm. (B) Quantitative analysis of the effect of PRG overexpression on actin polymerization at the Solo-accumulation site. The Solo-accumulation sites were extracted from the fluorescence images in A, and the intensities of actin fluorescence at these sites were measured and divided by the area. The relative intensity is shown as the mean ± SD of five independent experiments (32–59 areas/experiment), with the intensity of cells expressing mCherry and YFP-Solo set as 1. (C) Quantitative analysis of the effect of PRG and Solo overexpression on actin polymerization. The intensities of actin fluorescence in the basal area of the cells were measured and divided by the area of the cells. The region of each cell was specified by binarizing the fluorescence image of CFP-CAAX. The relative intensity is shown as the mean ± SD of five independent experiments (21–37 cells/experiment), with the intensity of cells expressing YFP and mCherry set as 1. (D) Quantitative analysis of the effect of Solo and PRG co-expression on PRG-induced lamellipodia formation. The number of cells with lamellipodia was counted from images of actin staining in A. The percentage of cells with lamellipodia to the total number of cells observed is shown as the mean ± SD of three independent experiments (22–39 cells/experiment). \**p* < 0.05, \*\**p* < 0.01, \*\*\**p* < 0.001, \*\*\*\**p* < 0.0001 (B, two-tailed paired *t*-test; C and D, one-way ANOVA followed by Tukey’s test).

Moreover, co-expression of Solo and PRG enhanced Solo-induced actin assembly on the basal plane (Figure 2A). To assess whether co-expression PRG with Solo affects actin filament assembly in Solo-expressing cells, we analyzed the fluorescence intensity of F-actin in the region of Solo accumulation on the basal plane to examine the effect of PRG expression on actin filaments. The relative intensity of actin filaments in the Solo-accumulated region was significantly higher in mCherry-PRG-co-expressing cells than in control mCherry-co-expressing cells (Figure 2B). These results suggest that PRG co-localizes with Solo and promotes Solo-induced actin filament assembly on the basal plane of the cell. Additionally, we analyzed the fluorescence intensity of F-actin in whole cells to investigate the effects of Solo co-expression on PRG-induced actin assembly. Solo co-expression with PRG increased the number and sickness of stress fibers and inhibited PRG overexpression-induced lamellipodia. Quantitative analysis revealed that Solo co-expression synergistically increased PRG-induced actin assembly and decreased PRG-induced lamellipodia formation (Figure 2, C and D).

### PDZ-RhoGEF knockdown suppresses Solo-induced actin assembly

To investigate the role of PRG in Solo localization and function, we examined the effects of PRG knockdown on Solo localization and Solo-induced actin filament assembly. MDCK cells were co-transfected with YFP or YFP-Solo expression plasmids and small interfering RNAs (siRNAs) targeting PRG. Immunoblot analysis showed that two independent PRG-targeting siRNAs effectively suppressed PRG protein expression in MDCK cells (Figure 3A). PRG knockdown suppressed stress fiber formation in YFP-expressing control cells (Figure 3B). Quantitative analysis of the fluorescence intensity of F-actin in the whole cell showed that actin assembly was significantly decreased in PRG-knockdown cells compared with that in control cells (Figure 3C). In cells expressing YFP-Solo, strong actin assembly was observed at the Solo-accumulation site; however, PRG knockdown suppressed Solo-induced actin assembly at the Solo-accumulation site (Figure 3D). Quantitative analysis of the fluorescence intensity of F-actin in the Solo-accumulated region revealed that Solo-induced actin assembly was significantly decreased in PRG-knockdown cells (Figure 3E). Collectively, these results suggest that Solo-induced actin assembly requires PRG. PRG knockdown had no significant effect on the linear localization of Solo in the basal plane, indicating that PRG was not involved in the localization of Solo in the basal plane.

**FIGURE 3.**
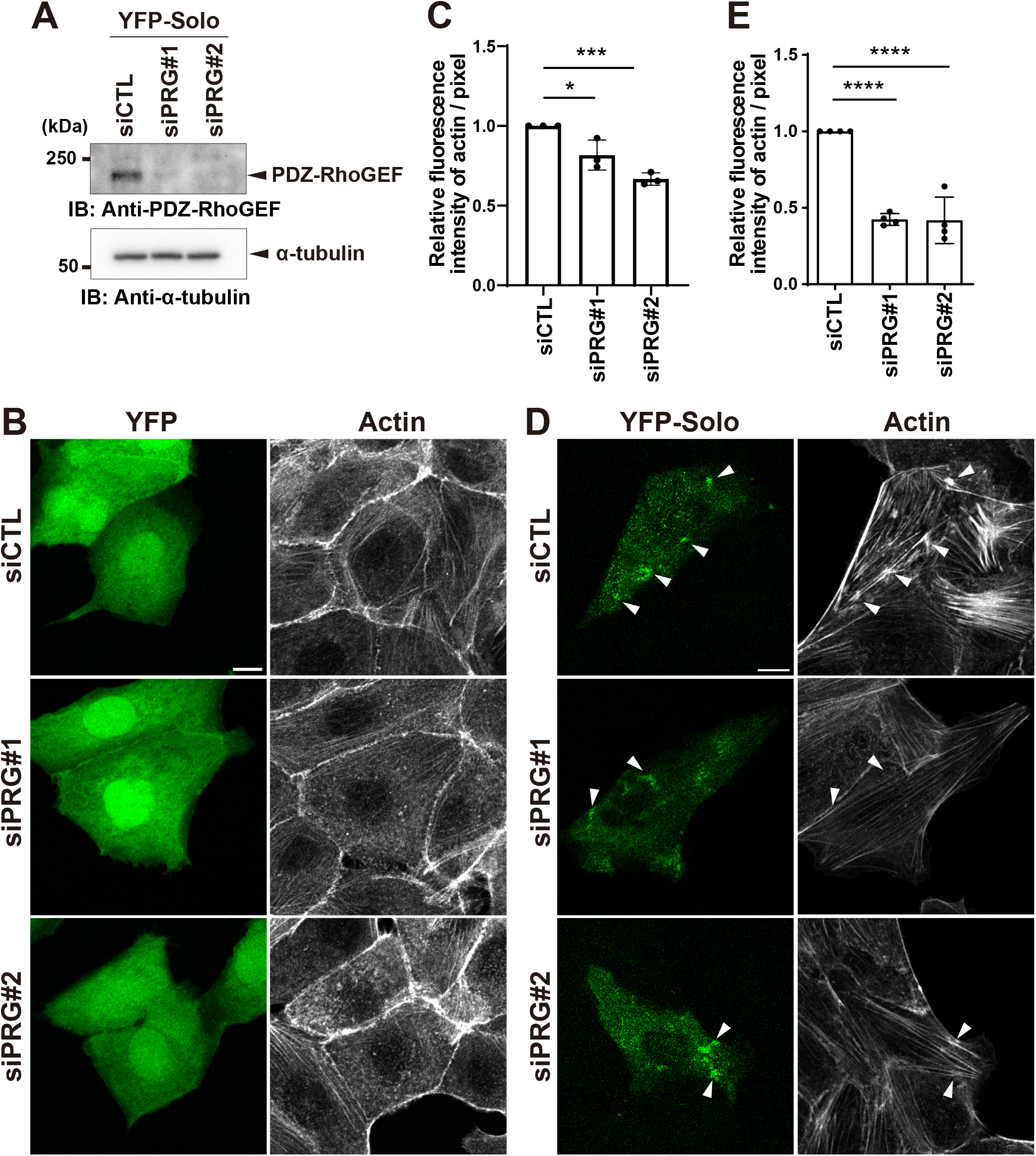
PRG knockdown suppresses Solo-induced actin polymerization. (A) Effects of siRNAs on Solo expression. MDCK cells were transfected with control siRNA or PRG-targeting siRNAs and cultured for 48 h. Cell lysates were analyzed by immunoblotting with anti-PRG antibody. (B) Confocal microscopic images of YFP and F-actin in the control and PRG knockdown cells. MDCK cells were transfected with YFP and CFP-CAAX expression plasmids and control siRNA or PRG-targeting siRNAs. (C) Quantitative analysis of the intensities of actin in the basal area of the cells as in Fig. 2B. The relative intensity is shown as the mean ± SD of three independent experiments (28–44 cells/experiment), with the intensity of control siRNA transfected cells set as 1. (D) Confocal microscopic images of YFP-Solo and F-actin in the control and PRG knockdown cells. MDCK cells were co-transfected with YFP-Solo expression plasmid and control siRNA or PRG-targeting siRNAs. White arrowheads indicate the accumulation sites of YFP-Solo. Scale bar, 10 µm. (E) Quantitative analysis of the effect of PRG knockdown on Solo-induced actin polymerization. The intensities of actin fluorescence in the Solo-accumulation sites were measured as in Fig. 2D. The relative intensity is shown as the mean ± SD of four independent experiments (38–68 areas/experiment), with the intensity of control siRNA transfected cells set as 1. \**p* < 0.05, \*\*\**p* < 0.001, \*\*\*\**p* < 0.0001 (one-way ANOVA followed by Dunnet’s test).

### Solo knockdown suppresses PDZ-RhoGEF-induced actin assembly

To elucidate the effect of Solo knockdown on PRG-induced actin filament assembly, MDCK cells were transfected with mCherry-PRG or siRNA-targeting Solo. Immunoblot analysis showed that Solo-targeting siRNAs reduced Solo protein expression in MDCK cells (Figure 4A). Solo knockdown in control mCherry-expressing cells slightly reduced stress fiber formation (Figure 4, B and C). In contrast, Solo knockdown in mCherry-PRG-expressing cells suppressed PRG-induced stress fiber formation on the basal plane, but not lamellipodium-like actin assembly at the cell periphery. Quantitative analysis of actin assembly in whole cells revealed that PRG expression increased actin assembly in the cells. On the other hand, Solo knockdown significantly suppressed PRG-induced actin assembly to the same level as that in control cells (Figure 4C).

**FIGURE 4.**
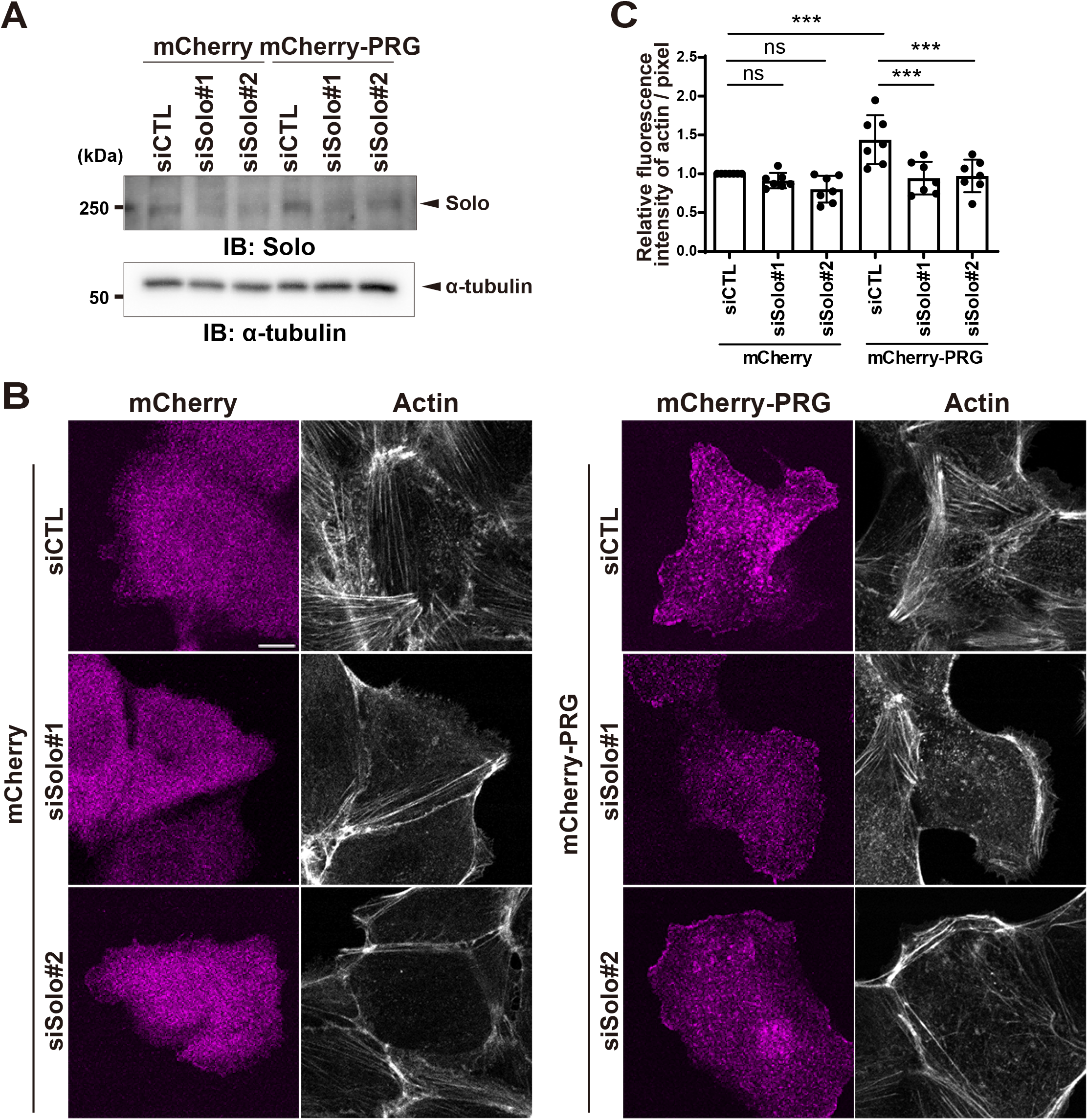
Solo knockdown partially suppresses PRG-induced actin polymerization. (A) Effects of siRNAs on Solo expression. MDCK cells were co-transfected with mCherry or mCherry-PRG, CFP-CAAX and control siRNA or Solo-targeting siRNAs and cultured for 48 h. Cell lysates were analyzed by immunoblotting with anti-Solo antibody. (B) Confocal microscopic images of mCherry or mCherry-PRG and F-actin in control and Solo knockdown cells. (C) Quantitative analysis of the intensities of actin in the basal area of the cells as in Fig. 2C. The relative intensity is shown as the mean ± SD of seven independent experiments (26–44 cells/experiment), with the intensity of mCherry and control siRNA transfected cells set as 1. \**p* < 0.05, \*\**p* < 0.01, \*\*\**p* < 0.001 (one-way ANOVA followed by Tukey’s test).

Overall, these results suggest that Solo promotes PRG-induced actin filament assembly. Therefore, it is possible that Solo and PDZ-RhoGEF act in synergy to stimulate actin filament assembly.

### Mapping of the binding regions of Solo and PRG

PRG contains an N-terminal PDZ domain, regulator of G protein signaling (RGS) homology domain, actin-binding domain (ABD), and tandem DH and PH domains (Figure 5A, Fukuhara *et al*., 1999; Banerjee *et al*., 2009). To determine the Solo-binding region of PRG, we constructed expression plasmids encoding mCherry-tagged wild-type (WT) PRG and its deletion mutants (Figure 5A). The plasmids were individually co-expressed with YFP-Solo in COS-7 cells, and their binding ability was analyzed via immunoprecipitation with an anti-GFP nanobody and anti-mCherry immunoblotting (Figure 5B). mCherry-PRG (WT) and its deletion mutant (N-170) co-precipitated with YFP-Solo, but the other deletion mutants did not, indicating that Solo binds to the N-terminal PDZ domain-containing region of PRG (Figure 5B). Solo contains an N-terminal Solo domain, followed by a CRAL/TRIO domain, spectrin repeats in the central region, and a C-terminal DH-PH domain (Figure 5C, Fujiwara *et al*., 2016). To map the PRG-binding region of Solo, YFP-tagged Solo (WT) and its deletion mutants (Figure 5C) were individually expressed in COS-7 cells, and their binding ability to endogenous PRG was analyzed via immunoprecipitation and immunoblotting (Figure 5D). Endogenous PRG co-precipitated with YFP-Solo (WT) and its deletion mutant (330-1057), but faintly with YFP-Solo (1058-C). Overall, these results indicate that Solo binds to PRG principally via the central (330–1057) region (Figure 5D).

**FIGURE 5.**
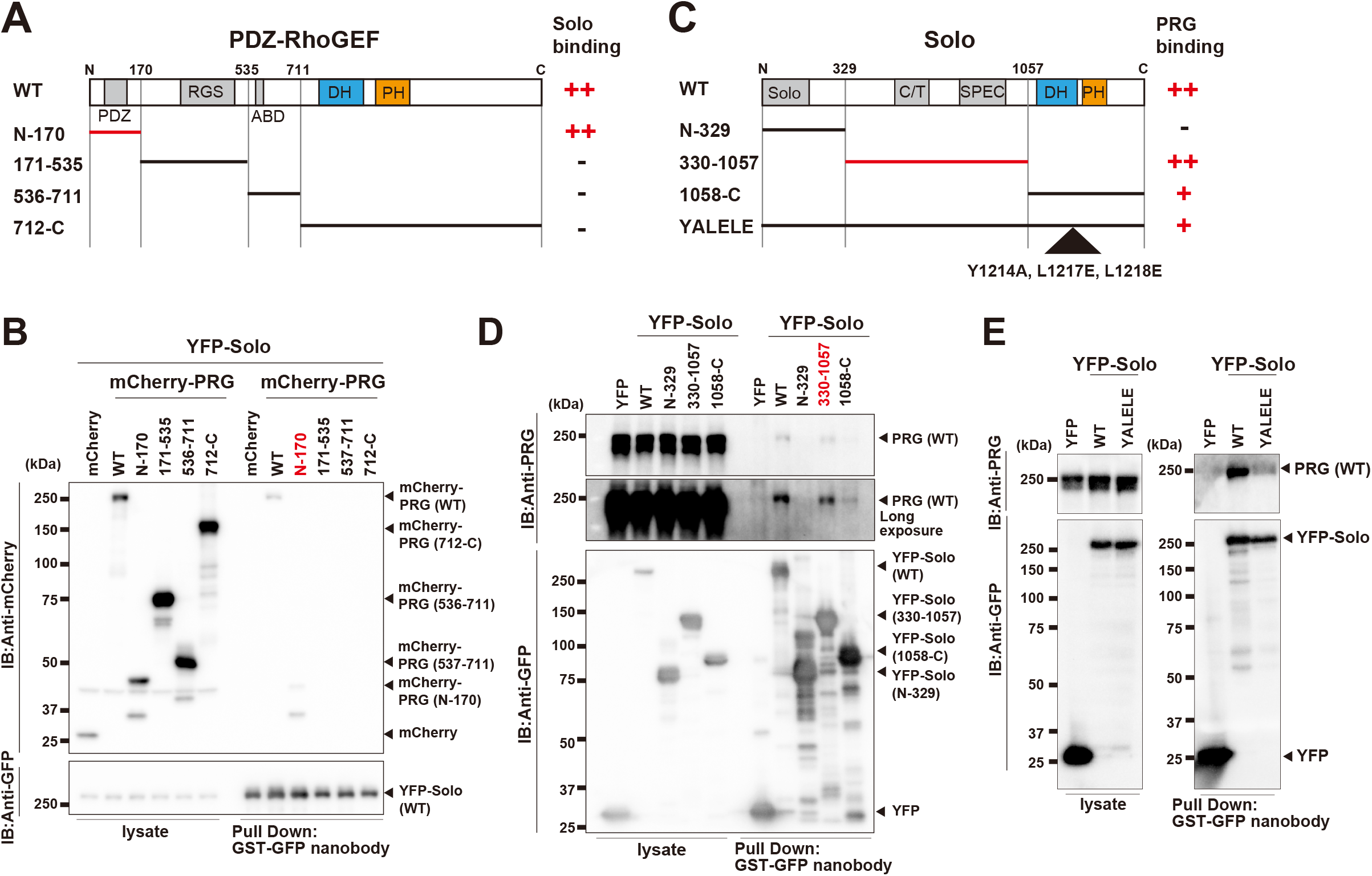
Determination of the binding region of Solo and PRG. (A) Schematic structures of PRG and its deletion mutants. PDZ: PDZ domain, RGS: regulator of G protein signaling domain, ABD: actin-binding domain (B) Identification of the Solo-binding region of PRG. Each deletion mutant of PRG was co-expressed with YFP-Solo in COS-7 and precipitated with GST-GFP nanobody. Precipitates were subjected to SDS-PAGE and analyzed by immunoblotting with anti-mCherry or -GFP antibodies. (C) Schematic structures of Solo and its deletion mutants. (D) Identification of the PRG-binding region of Solo. Each deletion mutant of Solo was precipitated with GST-GFP nanobody. Precipitates were analyzed by immunoblotting as in Figure 1D. The long exposure image of the blot with anti-PRG antibody is shown. (E) Co-immunoprecipitation assay of Solo inactive mutant and PRG. YFP, YFP-Solo, and YFP-Solo-YALELE inactive mutant were expressed in COS-7 cells and precipitated with GST-GFP nanobody. Precipitates were analyzed by immunoblotting as in Figure 1D.

Given that the localization of PRG is dependent on the site of Solo accumulation, we examined whether the association between Solo and PRG is dependent on the GEF activity of Solo. In a previous study, we showed that the L1217E mutant has a dominant-negative effect on mechanical stress responses, but it could not verify GEF activity. Therefore, we constructed a GEF-inactive Solo mutant (YALELE), in which Tyr-1214 was replaced by Ala and Leu-1217 and Leu-1218 were replaced by Glu in the DH domain, based on reports that β-Pix and GEF-H1 mutations can induce loss of GEF activity (Manser *et al*., 1998; Krendel *et al*., 2002). Co-precipitation assay revealed that YFP-Solo (YALELE) mutant co-precipitated endogenous PRG, but its binding affinity was extremely weak compared with that of Solo (WT) (Figure 5E).

### Solo-and PRG-induced actin polymerization requires Solo and PRG interaction

Furthermore, we examined the role of the interaction between Solo and PRG on their induced actin polymerization by overexpressing each deletion mutant as a dominant negative effect. MDCK cells were co-transfected with YFP-Solo and mCherry-tagged deletion mutants of PRG, and the effects on the actin cytoskeleton were quantitatively analyzed by fluorescence intensity of F-actin. Solo-induced actin polymerization at the Solo-accumulated sites was significantly suppressed following co-expression of mCherry-PRG (N-170) mutant (Figure 6, A and B). Although the PRG (712-C) mutant also significantly suppressed Solo-induced actin polymerization at the Solo-accumulated sites, this is probably because the DH-PH domain hyperactive mutant formed strong stress fibers, thereby depriving actin monomers. Similarly, co-expression of mCherry-PRG and YFP-Solo (330–1057) mutant in MDCK cells significantly suppressed PRG-induced actin polymerization (Figure 6, C and D). The putative GEF-inactive mutant of Solo (YALELE) did not accumulate at the basal side of the cells and did not inhibit PRG-induced actin polymerization. Co-precipitation analysis revealed that Solo mutation reduced its interaction with PRG (Figure 5E), indicating that an interaction between Solo and PRG is required for PRG-induced actin stress fiber formation. Collectively, these results suggest that the interaction between Solo and PRG is required for the RhoGEFs to exert their specific actin cytoskeletal remodeling.

**FIGURE 6.**
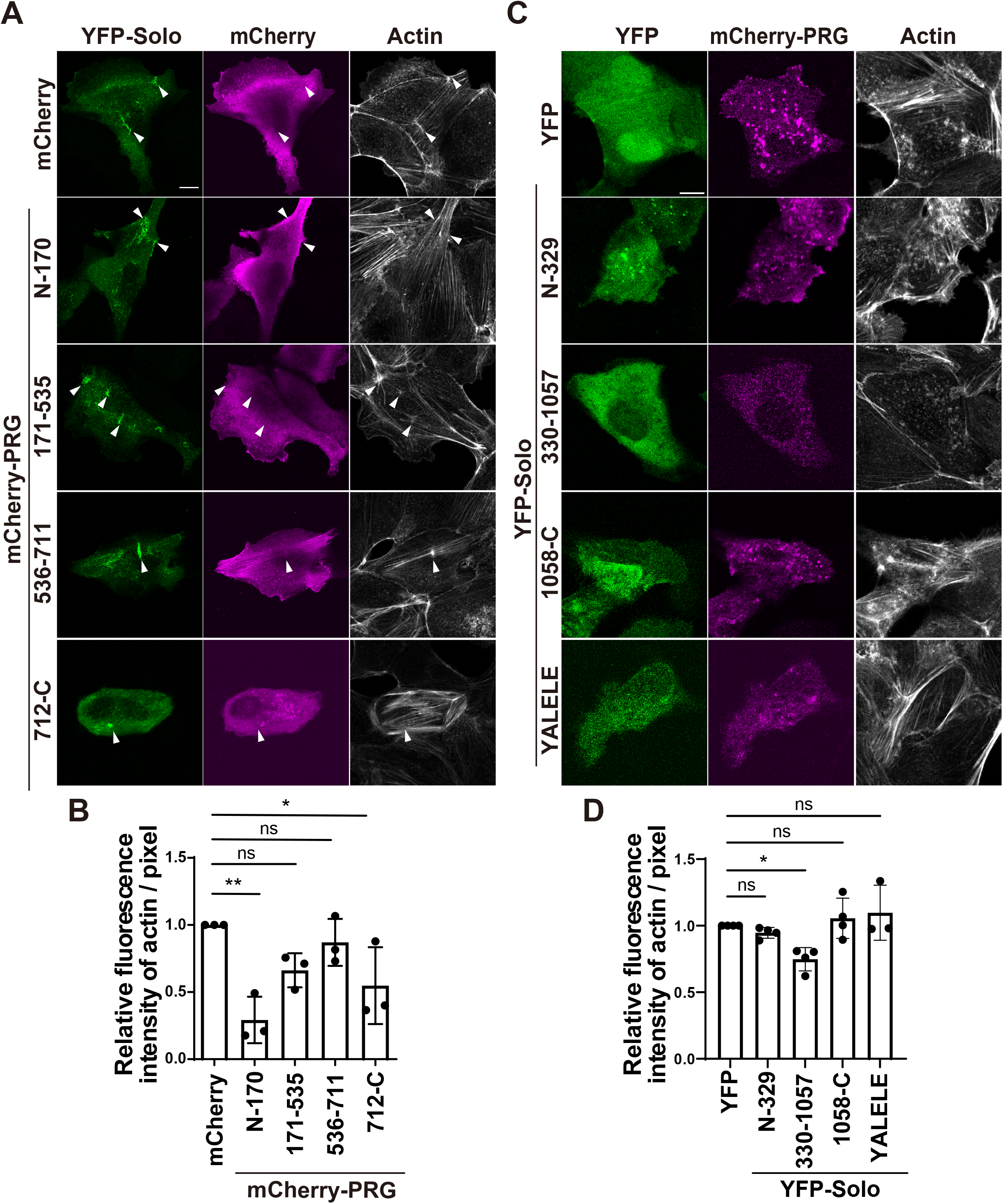
Solo or PRG-induced actin polymerization is suppressed by the overexpression of the Solo-binding region of PRG or PRG-binding regions of Solo. Overexpression of the Solo or PRG-binding regions suppresses actin polymerization induced by the proteins. (A, B) Confocal microscopic images of YFP, YFP-Solo, or its mutants; mCherry, mCherry-PRG, or its mutants; and F-actin in MDCK cells. MDCK cells were co-transfected with YFP-Solo, mCherry, or each mCherry-PRG deletion mutant and CFP-CAAX in A, and co-transfected with mCherry-PRG and YFP, each YFP-Solo deletion mutant, or an inactive mutant of Solo and CFP-CAAX in B. Cells were fixed and stained as in Figure 2A. White arrowheads in A indicate the accumulation sites of YFP-Solo. Scale bar, 10 µm. (C) Quantitative analysis of intensity of F-actin at Solo accumulation sites. The relative intensity is shown as the mean ± SD of three independent experiments (29–61 areas/experiment), with the intensity of mCherry and YFP-Solo transfected cells set as 1. (D) Quantitative analysis of intensity of F-actin in the basal area of the cells. The relative intensity is shown as the mean ± SD of three or more independent experiments (22–36 cells/experiment), with the intensity of cells expressing YFP and mCherry-PRG set as 1. \**p* < 0.05, \*\**p* < 0.01 (one-way ANOVA followed by Dunnett’s test).

### Solo- and PRG-induced RhoA activation requires Solo and PRG interaction

Both Solo and PRG are RhoGEFs that target RhoA and induce stress fiber formation. In the present study, PRG enhanced Solo-induced actin polymerization at the site of Solo accumulation, indicating that Solo and PRG interaction may affect their GEF activity. Therefore, we examined the effect of PRG knockdown on Solo-induced RhoA activation using an active RhoA pulldown assay. Two independent MDCK cell lines constitutively expressing YFP-Solo were used for experiment because the transfection efficiency was low, and Solo-induced RhoA activation was too weak to detect the effect of PRG knockdown on RhoA activity. The RhoA activities of the two YFP-Solo-expressing cell lines were higher and comparable to that of YFP-expressing MDCK cell lines as using control cells (Figure 7A, B). The lack of an increase in RhoA activity in the YFP-Solo-expressing cell line may be because other RhoA GEFs were suppressed by the homeostatic mechanism of RhoA activity during the establishment of the cell line. Therefore, PRG knockdown significantly reduced RhoA activity in both YFP- and YFP-Solo-expressing cells, with YFP-Solo-expressing cells exhibiting a greater decrease than YFP-expressing cells (Figure 7, C and D). Furthermore, we examined the effect of Solo-knockdown on PRG-induced RhoA activation. Solo was knocked down in MDCK cells transiently expressing PRG, and RhoA activity was measured. Transient expression of PRG strongly activated RhoA in MDCK cells; however, Solo knockdown suppressed RhoA activity in control cells and even more strongly in PRG-expressing cells (Figure 7, E–H). Overall, these results suggest that Solo and PRG activate each other to activate RhoA.

**FIGURE 7.**
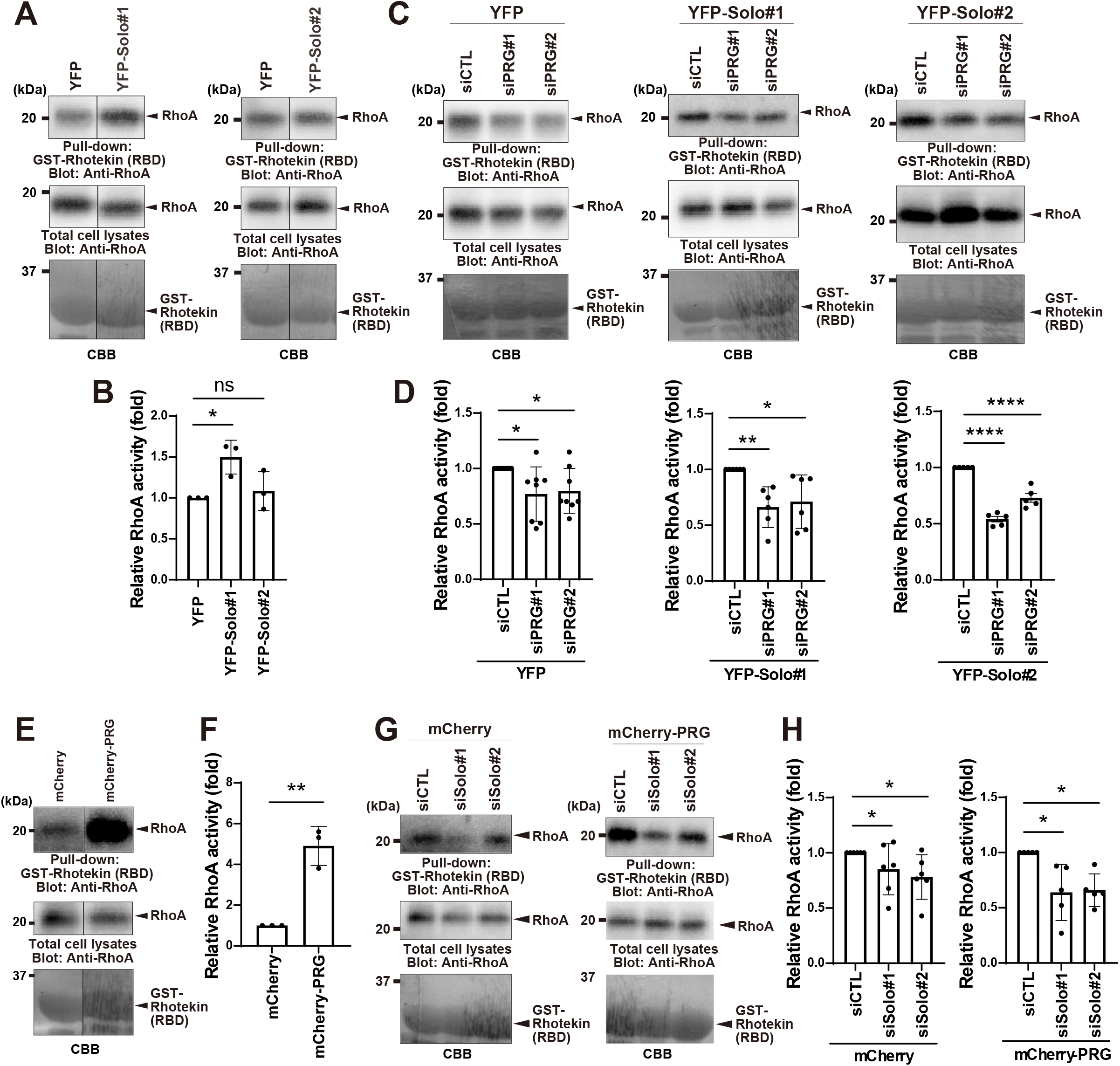
Interaction between Solo and PRG is involved in RhoA activation by each of them (A) RhoA activities in MDCK cells and Solo-expressing MDCK cell lines. Cells of the parental MDCK cell line and two independent Solo-expressing MDCK cell lines were lysed and subjected to a GST-Rhotekin (RBD) pulldown assay. Total cell lysates and precipitates were analyzed by immunoblotting with anti-RhoA antibody. (B) Quantitative analysis of relative RhoA activity is shown as the mean ± SD of three or more independent experiments, with the activity in YFP-expression MDCK cells set as 1. (C) The effect of PRG knockdown on Solo-mediated RhoA activation. MDCK cells stably expressing YFP or YFP-Solo were transfected with control and PRG-targeting siRNAs. RhoA activity was measured using a GST-RBD pulldown assay as in A. (D) Quantitative analysis of relative RhoA activity is shown as the mean ± SD of three or more independent experiments, with the activity in control siRNA transfected cells set as 1. (E) Effect of PRG overexpression on RhoA activation. RhoA activity in MDCK cells expressing mCherry or mCherry-PRG was measured using a GST-RBD pulldown assay as in A. (F) Quantitative analysis of relative RhoA activity is shown as the mean ± SD of five or more independent experiments, with the activity in mCherry-expression MDCK cells set as 1. (G) The effect of Solo knockdown on PRG-induced RhoA activation. MDCK cells were co-transfected with mCherry or mCherry-PRG expression plasmid and control siRNA or Solo targeting siRNAs. RhoA activity was measured using a GST-RBD pulldown assay. (H) Quantitative analysis of relative RhoA activity is shown as the mean ± SD of five or more independent experiments, with the activity in control siRNA transfected cells set as 1. (B and F, two-tailed paired *t*-test; D and H, one-way ANOVA followed by Dunnett’s test) \**p* < 0.05, \*\**p* < 0.01, \*\*\*\**p* < 0.0001.

### Interaction between Solo and PRG is required for actin cytoskeletal remodeling in response to substrate stiffness

Solo and PRG are involved in mechanical stress response in cells (Fujiwara *et al*., 2016; Ito *et al*., 2017), co-localized to the basal area, and induce stress fiber formation, indicating that the interaction is involved in the response of cells to substrate stiffness. Therefore, we investigated whether the interaction between Solo and PRG is involved in actin cytoskeletal remodeling in response to substrate stiffness. Given that cell shapes are spread to nearly the same size, the number of stress fibers is very low at 10 kPa, and as many as above 35 kPa. Therefore, we measured the difference in the number of stress fibers on the gel at 10 and 35 kPa to examine the effects of substrate stiffness on changes in actin cytoskeleton in cells transfected with the deletion mutants of Solo and PRG, which are binding regions of each other. The number of YFP-expressing control cells with more than five stress fibers was significantly higher in the 35 kPa substrate group than in the 10 kPa substrate group. In contrast, there was no significant difference in the number of cells with more than five stress fibers between the 35 and 10 kPa substrate groups, expressing with the Solo (330–1057) or PRG (N-170) mutants (Figure 8, A and B). Additionally, Solo/PRG knockdown inhibited actin polymerization, including stress fiber formation, in cells seeded on glass (3Figure 3B and 4B). Overall, these results suggest that the interaction between Solo and PRG is required for actin cytoskeletal remodeling in response to substrate stiffness. This signaling cascade may be important for the adaptation of cells to the extracellular mechanical environment.

**FIGURE 8.**
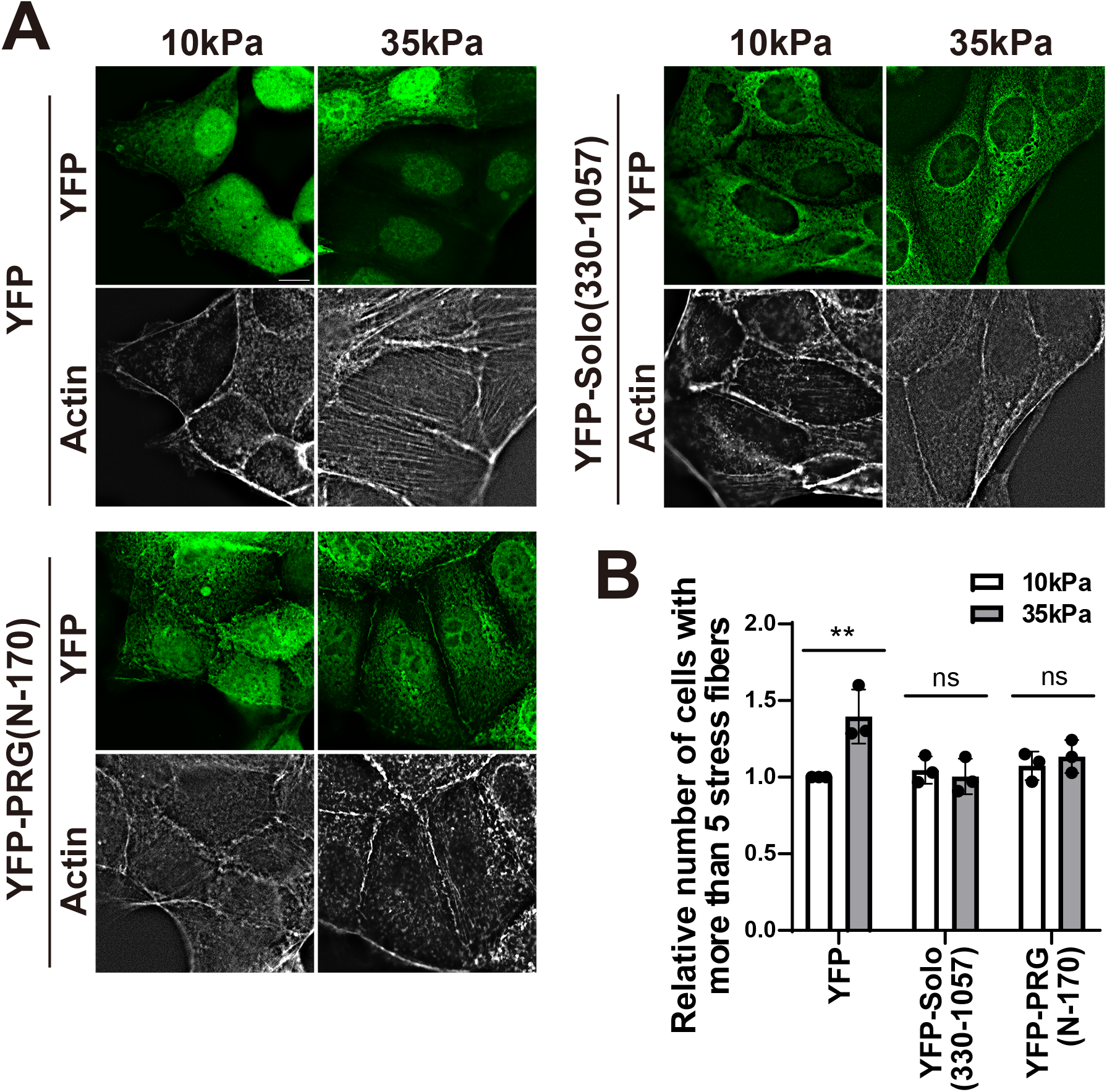
Solo and PRG interaction is required for substrate stiffness-dependent stress fiber formation in MDCK cells. (A) Fluorescence images with deconvolution of YFP, YFP-Solo (330– 1057), or YFP-PRG(N-170) and F-actin in MDCK cells on the 10 kPa or 35 kPa fibronectin-conjugated PA-gel. Scale bar, 10 µm. (B) Quantitative analysis of the number of cells with more than five stress fibers. The relative number is shown as the mean ± SD of three independent experiments (27–99 cells/experiment), with the intensity of cells expressing YFP set as 1. \*\**p* < 0.01 (Two-Way ANOVA followed by Sidak’s test)

## DISCUSSION

In this study, we identified various candidate Solo-interacting proteins by proteomic analysis using BioID. Among the interacting proteins identified, PRG interacted with Solo and cooperatively reorganized the characteristic actin cytoskeletal structure, and this interaction was involved in cell response to substrate stiffness. Other candidates include proteins involved in regulating actin cytoskeleton and intermediate filaments, proteins that support the membrane cytoskeleton, and cell-cell and cell-substrate adhesion. Solo is involved in mechanical stress response and localizes in punctate dots in the basal area of cells (Fujiwara *et al*., 2016). These findings suggest that Solo is associated with membrane cytoskeletal proteins, such as spectrin and myosin I, which are involved in mechanical stress response. However, detailed analyses are necessary to determine their relationship with Solo.

In the present study, PRG and LARG (members of the RGS-RhoGEF family) were identified as Solo-interacting proteins. A comprehensive proteomic analysis of RhoGEF and LARG using the BioID method has confirmed these interactions with Solo (O’Loughlin *et al*., 2018; Muller *et al*., 2020). LARG and PRG are involved in tensile force-induced RhoA activation and force regulation of cell-cell adhesion sites, respectively (Guilluy *et al*., 2011b; Ito *et al*., 2017), suggesting that the interaction between PRG or LARG and Solo may be involved in mechanical stress responses. Co-expression analysis of Solo and PRG showed that PRG changed localization from the cell periphery to the Solo accumulation site in the cell interior depending on the localization of Solo. Additionally, PRG-induced lamellipodia formation was suppressed and ventral stress fibers in the cell interior were strongly induced (Figure 2). Overall, these results suggest that Solo regulates the target specificity of PRG and the intracellular sites where PRG is activated. The effects of PRG overexpression on stress fiber formation and RhoA activation tended to be much stronger than those of Solo overexpression (Figure 2). PRG activates RhoA and Cdc42 downstream of G protein-coupled receptors (GPCR) through the interaction between the trimeric G protein and the RGS domain, resulting in the induction of lamellipodia and stress fiber formation in the peripheral region (Driessens *et al*., 2002; Banerjee and Wedegaertner, 2004). Solo knockdown does not suppress GPCR-induced activation of RhoA by lysophosphatidic acid (LPA), but it suppresses tensile force-induced RhoA activation (Fujiwara *et al*., 2016). These findings suggest that Solo-dependent PRG activation is independent of the GPCR signaling pathway. PRG is a RhoGEF that activates RhoA and Cdc42 downstream of multiple signaling pathways to form substantial actin structures, and Solo promotes PRG-mediated RhoA activation in response to the external mechanical environment. However, Solo-or PRG-induced characteristic actin cytoskeletal remodeling and activation of RhoA were suppressed in cells overexpressing Solo or PRG by the knockdown of other RhoGEFs (Figure 3, 4, and 7). These results indicate that Solo and PRG do not engage in a cascade but rather act synergistically.

Although little is known about the activation mechanism of Rho GTPases, which acts in a feed-forward manner by binding between RhoGEFs, it has been reported that RhoGEFs negatively regulate the activity of other RhoGEFs and RhoGAPs via Rho GTPases or direct binding (Guilluy *et al*., 2011a; Muller *et al*., 2020). For example, the binding of Cdc42 to α-Pix activates the GEF activity of α-Pix for Rac, and β-Pix (a Rac/Cdc42-GEF) forms a complex with srGAP1 (a RhoA-GAP) to regulate the activity of Rho GTPases (Baird *et al*., 2005; Kutys and Yamada, 2014). Additionally, BPGAP, a GAP of RhoA, binds to Vav1 in lamellipodia, inactivates RhoA, and facilitates the GEF activity of Vav1 toward Rac, resulting in the promotion of lamellipodia formation (Wong *et al*., 2023). Moreover, PLEKHG4B, which has a structure similar with Solo, binds to PDZ-RhoGEF and LARG and inhibits the activation of their GEF activities downstream of the G-protein coupled receptor (Muller *et al*., 2020; Ninomiya *et al*., 2021). Conclusively, these findings suggest that the activity of Rho GTPases is diversely regulated by interactions with upstream regulators.

PRG, LARG, and p115RhoGEF belong to the RGS-RhoGEF family with an RGS domain. RGS-RhoGEF is activated downstream of GPCRs and is involved in force-induced RhoA activation (Kozasa *et al*., 2011; Lessey *et al*., 2012). In the present study, we found that the PDZ domain at the N-terminus of PRG is the binding domain of Solo. LARG and PRG, which contain PDZ domains, interact with Solo (O’Loughlin *et al*., 2018; Muller *et al*., 2020). Therefore, the PDZ domains of PRG and LARG are responsible for the Solo-mediated signaling pathway. Research findings indicate that Solo is required for the activation of RhoA by mechanical forces through VE-cadherin and integrin (Abiko *et al*., 2015; Fujiwara *et al*., 2016). Moreover, Solo binds to keratin 8/keratin 18 (K8/K18) intermediate filaments, and this interaction is required for force-induced RhoA activation (Fujiwara *et al*., 2016). PRG is involved in the regulation of forces at cell-cell adhesion sites in epithelial cells, and LARG is required for the regulation of contractile forces through cell-substrate adhesion sites (Guilluy *et al*., 2011b; Ito *et al*., 2017). In the present study, we showed that the interaction between Solo and PRG is involved in the cellular response to substrate stiffness, a mechanical stress response. Several molecular mechanisms have been reported for this response to substrate stiffness. Integrins form focal adhesions (FA) link actomyosin via talin in the cytoplasmic domain. The contractile force generated by actomyosin is then subjected to talin, counteracting the stiffness of the substrate, which leads to conformational changes in talin, resulting in talin functioning as a stiffness sensor (Elosegui-Artola *et al*., 2016). The mechanosensory mechanism of integrin involves the activation of RhoA via focal adhesion kinase (FAK) and Src activation (Guilluy *et al*., 2011b). Upstream of RhoA, p190RhoGEF, p115RhoGEF, and p190RhoGAP regulate RhoA activity through mechanosensory signals at the FA (Lessey *et al*., 2012). The functions of Solo and PRG may depend on these stiffness-sensing mechanisms. Therefore, it could be speculated that Solo binds to K8/K18, an intermediate filament, and anchors keratin fibers to the membrane of the basal area of cells (Fujiwara *et al*., 2016). These findings suggest a novel molecular mechanism by which the tensile force from cell-substrate adhesions accumulates and activates Solo in the vicinity via keratin fibers. Solo then accumulates PRG and generates a contractile force in response to the tensile force applied to the cell caused by the substrate stiffness.

To conclude, Solo dominantly regulates the localization of PRG, and the interaction between Solo and PRG is involved in reorganizing the actin cytoskeleton in response to substrate stiffness. However, the localization of Solo in the basal area of cells has not been well characterized, and the function of keratin IFs networks in response to substrate stiffness remains unknown. Therefore, further studies are necessary to comprehensively elucidate the molecular mechanisms of Solo localization and activation.

## MATERIALS AND METHODS

### Reagents and antibodies

Alexa 647-phalloidin was purchased from Thermo Fisher Scientific (A30107). The following antibodies were used for the study: mouse anti-GFP (632381, JL-8, Clontech, 1:1000), rabbit anti-mCherry (GTX128508, Gene Tex, 1:1000 for WB, 1:500 for IF), rabbit anti-PDZ-RhoGEF (ab110059, Abcam, 1:1000), mouse anti-RhoA (sc-418, 26C4, Santa Cruz Biotechnology, 1:250), mouse anti-α tubulin (T5168, B-5-1-2, Sigma-Aldrich, 1:1000). Rabbit anti-Solo antibody was purified using an immunogen peptide conjugated-Sepharose column from the antiserum, which was raised against the C-terminal peptide (LSRQSHARALSDPTTPL) of human Solo (Abiko *et al*., 2015). Alexa 488-conjugated anti-mouse IgG (A28175, 1:500), Alexa 568-conjugated goat anti-rabbit IgG (A11036, 1:500), horseradish peroxidase (HRP)-conjugated anti-mouse IgG (NA931, 1:10000), and anti-rabbit IgG (NA934, 1:10000) were purchased from Thermo Fisher Scientific.

### siRNAs

The following siRNAs were purchased from Sigma-Aldrich for the experiments: 5’-GAGCUGAAAGAGGAACUCAAACC-3’ (dog *Solo* #1), 5’-GGGAUCAGAGACCUUUGUUUACA-3’ (dog *Solo* #2), 5’-GAGCTCATAGAGATCCACA-3’ (dog *PDZ-RhoGEF* #1), and 5’-GGGAAAUUCUCAAGUACGU-3*’* (dog *PDZ-RhoGEF* #2). The negative control siRNA was purchased from Sigma-Aldrich (MISSION siRNA Universal Negative Control, SIC001).

### Plasmid construction

A plasmid containing the cDNA encoding TurboID was purchased from Addgene (107169). The pCAG-V5-TurboID vector was constructed by inserting the amplified TurboID cDNA containing the V5 epitope (GKPIPNPLLGLDST) sequence at the N-and C-terminal sites into the pCAGGS expression plasmid (Addgene). Expression plasmids encoding V5-TurboID-Solo and Solo-TurboID-V5 were constructed by inserting amplified Solo cDNA with a GS-linker (GGGSx2) sequence at the N-and C-terminal sites, and TurboID cDNA with a V5 epitope tag at the N-and C-terminal sites, respectively, into the pCAGGS vector. Human PDZ-RhoGEF cDNA was amplified using PCR with cDNA derived from cultured human smooth muscle (SMC) cells and the following primers: forward (fw) 5′-ACAAGTCTCGAGGGTACCAGAGTGTAAGGTTACCCCAGAG-3′ and reverse (rv) 5′-GATATCAAGCTTGTCGACTTATGGTCCTGGTGACGCGG-3. Expression plasmids encoding mCherry-tagged PDZ-RhoGEF and its mutants were constructed by inserting the PCR-amplified cDNA fragments into an mCherry-C1 expression plasmid (Clontech). The cDNA fragments were amplified using PrimeSTAR HS DNA polymerase (TAKARA, Japan), and plasmid recombination was performed using NEBuilder HiFi DNA assembly (New England Biolabs). The expression plasmid encoding CFP-CAAX was constructed by inserting the CAAX sequence (KDGKKKKKKSKTKCVIM), derived from the C-terminal sequence of human K-Ras, into ECFP-C1 expression plasmid (Clontech).

### Cell culture and transfection

DLD-1, MDCK, and COS-7 cells were obtained from the Riken BRD (Japan) and cultured in D-MEM (Wako, Japan) supplemented with 10% fetal bovine serum (FBS), 4 U/ml penicillin, and 40 µg/ml streptomycin at 37℃ in a 5% CO_2_ condition. Plasmid DNAs was transfected into DLD-1, MDCK, and COS-7 cells using Lipofectamine 2000 (Thermo Fisher Scientific), Lipofectamine LTX (Thermo Fisher Scientific), and JetPEI (Polyplus, France), respectively. Cultured cells were transfected with siRNAs using Lipofectamine RNAiMAX (Thermo Fisher Scientific). For immunofluorescence assay, mixtures of plasmid DNA/Lipofectamine LTX and siRNA/Lipofectamine RNAiMAX were added to the cells. For Rhotekin pull-down assay, a mixture of plasmid DNA and siRNA/Lipofectamine 2000 was added to the cells. DLD-1 cells constitutively expressing V5-TurboID, Solo-TurboID-V5, and V5-TurboID-Solo were established via puromycin selection (3.5 µg/ml). Cell lines were checked for mycoplasma every 3 months.

### Proteomic analysis

DLD-1 cells stably expressing V5-TurboID, V5-TurboID-Solo, Solo-TurboID-V5, and the parental DLD-1 cells were cultured in a medium containing biotin (50 mM) for 4 h. Thereafter, the cells were lysed with lysis buffer (25 mM Tris-HCl [pH 7.5], 140 mM NaCl, 2.5 mM MgCl_2_, 1 mM EDTA, 1% TritonX-100, 0.1% deoxycholic acid, and 1 mM DTT) and 1% Protease Inhibitor Cocktail (General Use, Nacalai Tesque, Japan), followed by centrifugation at 100,000 × g for 30 min at 4℃. DNase I (10 µg/ml) and RNase A (10 µg/ml) were added to the cell supernatants, followed by incubation with tamavidin-conjugated magnetic beads (MagCapture™ Tamavidin^®^2-REV, Wako) at 4℃ for 4 h. After washing the beads five times with wash buffer (25 mM Tris-HCl [pH 7.5], 140 mM NaCl, 2.5 mM MgCl_2_, 1 mM EDTA, 0.1% TritonX-100, and 1 mM DTT), biotinylated proteins were eluted with elution buffer (25 mM Tris-HCl [pH, 7.5], 0.5 M NaCl, and 20 mM biotin) twice at 4℃, 30 min first time and overnight the second time. The eluted proteins were subjected to acetone precipitation and dissolved in an SDS sample buffer (Nacalai Tesque). Proteins were separated using SDS-PAGE and detected using the Silver Stain MS Kit (Wako). Protein bands were excised and proteins in the gels were trypsinized. The digested peptides were recovered with a ZipTip C18 column (Millipore) and analyzed using matrix-assisted laser desorption/ionization time-of-flight tandem mass spectrometry (TOF/TOF 5800 system; AB Sciex, Framingham, MA, USA). Proteins were identified using MS-Fit software (http://prospector.ucsf.edu/prospector/mshome.htm) for MS data, and ProteinPilot software v3.0 (AB Sciex) for MS/MS data.

### Immunofluorescence staining and fluorescence imaging

Briefly, cells were cultured on coverslips, transfected with DNA plasmids or siRNAs, fixed with 4% paraformaldehyde at room temperature for 20 min, and permeabilized with 0.5% TritonX-100 at room temperature for 20 min. After blocked with 5% FBS in PBS, cells were incubated with primary antibodies diluted with Can Get Signal^®^ immunostain Solution A (TOYOBO, Japan) overnight at 4℃. After washing with PBS, the cells were incubated with secondary antibodies diluted with 25 mM Tris-HCl (pH, 7.4), 150 mM NaCl, 0.05% Tween20, and 1 mg/ml BSA at room temperature for 1 h, and then washed with PBS. Fluorescence images were obtained using an LSM710 laser-scanning confocal microscope (Carl Zeiss) equipped with a PL Apo 63× or 100× oil objective lens (1.4 NA), a DMI 8 fluorescence microscope (Leica Microsystems) equipped with a PL Apo 63× or 100× oil objective lens (1.4 NA), and an ORCA-Flash4.0 Digital CMOS camera (HAMAMATSU Photonics, Japan). Fluorescence images of the cells on polyacrylamide gels obtained using DMI 8 were improved using a 3D non-blind deconvolution method in the LAS X software (Leica Microsystems).

### Measurement of actin polymerization

Actin polymerization at the Solo accumulation sites and in the total basal area of cells was measured by assessing the fluorescence intensity of Alexa 647-phalloidin staining for F-actin using the ImageJ software (NIH). To measure the intensity of F-actin at the Solo accumulation sites, confocal images of the basal plane of YFP-Solo-expressing cells were binarized to segment the Solo-accumulated sites, and F-actin intensities in the segmented area were measured. To measure the intensity of F-actin in the total basal area of cells, cell shape was visualized based on the expression of plasma membrane-targeting signal-(K-Ras-CAAX)-tagged cyan fluorescent protein (CFP). Confocal images of the basal plane of CFP-CAAX-expressing cells were binarized to segment the total basal area of the cells, and the intensity of F-actin in this area was measured. The degree of F-actin polymerization was expressed as the average pixel intensity.

### Protein expression in *E*. *coli* and purification of GST-GFP nanobody

GST-GFP nanobody-expressing BL21 (DE3) cells were cultured at 37℃ until OD600 reached 0.5, followed by the addition of IPTG (0.1 mM) and further incubation for 4 h at 30°C. *E*. *coli* pellets obtained from 50 ml of culture were suspended in 10 ml of binding buffer (5 mM DTT, protease inhibitor cocktail in PBS) on ice and sonicated. Cell lysate was incubated with TritonX-100 for 30 min, followed by centrifugation at 20700 × g for 20 min at 4℃. Thereafter, the supernatant was incubated with 1500 µl of glutathione-Sepharose beads (50% slurry, GE Healthcare) at 4℃ for 4 h. The GST-GFP nanobody-bound Sepharose beads were washed with wash buffer (5 mM DTT, 0.1% TritonX-100 in PBS) and stored at 4℃.

### Protein expression in *E*. *coli* and purification of GST-Rhotekin-RBD

GST-Rhotekin RBD-expressing BL21 (DE3) cells were cultured at 37℃ until OD600 reached 0.5, followed by the addition of IPTG (0.5 mM) and incubation for 2 h. Thereafter, *E*. *coli* pellets obtained from 400 ml of cell culture were suspended in 10 ml of lysis buffer (50 mM Tris-HCl [pH, 7.4], 1% TritonX-100, 150 mM NaCl, 5 mM MgCl_2_, 1 mM DTT, 2 µg/ml pepstatin A, 10 µg/ml leupeptin, and 1 mM PMSF) on ice and sonicated. Cell lysate was centrifuged at 12000 × g for 20 min at 4℃ to collect the supernatant. Thereafter, cell supernatant was incubated with 500 µl of glutathione-Sepharose beads (50% slurry, GE Healthcare) at 4℃ for 1 h. Finally, GST-Rhotekin RBD-bound Sepharose beads were washed with wash buffer (50 mM Tris-HCl [pH, 7.4], 0.5% TritonX-100, 150 mM NaCl, 5 mM MgCl_2_, 1 mM DTT, 1 µg/ml pepstatin A, 5 µg/ml leupeptin, and 0.5 mM PMSF) and resuspended in 10% glycerol containing wash buffer and stored at –80℃.

### Co-precipitation assay

COS-7 cells (1.0 × 10^6^) in a 10-cm dish were transfected with plasmids encoding YFP-or mCherry-tagged proteins. Thereafter, the cells were washed with PBS, scraped, and collected in 1.5 ml tubes, followed by centrifugation at 8400 × g for 3 min at 4℃. Cell pellets were suspended and lysed with 0.5 ml of lysis buffer per dish (25 mM Tris-HCl [pH, 7.4], 140 mM NaCl, 1% TritonX-100, 2.5 mM MgCl_2_, 1 mM EGTA, 2 µg/ml pepstatin A, 10 µg/ml leupeptin, and 250 µM PMSF) and incubated for 5 min on ice. Cell lysate was centrifuged at 18800 × g for 10 min at 4℃, and the supernatants were incubated with GST-GFP nanobody-bound glutathione-Sepharose beads (GFP-nanobody Sepharose) at 4℃ for 4 h (Harmsen and De Haard, 2007). Finally, the GFP-nanobody Sepharose beads were washed with lysis buffer and proteins were eluted by boiling in Laemmli sample buffer (50 mM Tris-HCl [pH, 6.8], 1.6% SDS, 8% glycerol, 20% 2-mercaptoethanol, and 0.04% BPB) to elute the precipitated proteins for 3 min. The eluted proteins were separated using SDS-PAGE and analyzed using western blotting.

### Active RhoA pulldown assay

RhoA activity was evaluated by pulldown of its GTP-bound form using the GST-tagged RhoA-binding domain (RBD) of rhotekin, as previously described (Nishita *et al*., 2002). MDCK cells (3.0 × 10^5^) in 6 cm dishes were transfected with plasmids or siRNAs. After serum-starvation for 18 h, cells were washed with PBS and lysed with lysis buffer (50 mM Tris-HCl [pH, 7.4], 500 mM NaCl, 1% TritonX-100, 0.5% Na-deoxycholate, 0.1% SDS, 10 mM MgCl_2_, 1 mM DTT, 200 µM Na_3_VO_4_, 2 µg/ml pepstatin A, 10 µg/µl leupeptin, and 1 mM PMSF). Thereafter, cell lysate was centrifuged at 18800 × g for 5 min at 4℃, and the supernatant was incubated with GST-Rhotekin RBD-bound Sepharose beads for 1 h at 4℃. Proteins were eluted using Laemmli sample buffer and detected using western blotting. Relative RhoA activity was calculated by dividing the signal intensity of active RhoA by that of total RhoA. The intensity of the band signals was quantified using the ImageJ plugin and band/peak quantification tool (Ohgane and Yoshioka, 2019).

### Western blotting

Proteins were separated using sodium dodecyl sulfate-polyacrylamide gel electrophoresis and transferred to a polyvinylidene difluoride (PVDF) membrane (Immobilon, MILLIPORE). Membranes were blocked with 5% nonfat dry milk in PBS with 0.05% Tween20 (PBS-T) at room temperature for 1 h. To detect proteins, the membranes were subjected to immunoblot analysis as previously described (Ohashi *et al*., 2000). Chemiluminescence signals were generated using Immobilon Western Chemiluminescent HRP Substrate (Millipore) and measured using a Chemidoc Touch (Bio-Rad).

### Polyacrylamide hydrogels fabrication

Coverslips were hydrophilized with 1 M NaOH for 30 min at room temperature. After removing NaOH, the coverslips were incubated with 16.7% 3-aminopropyltrimethoxysilane in 1 M NaOH for 15 min at room temperature for silane coating. The coverslips were washed with water, incubated with 0.5% glutaraldehyde in PBS for 30 min at room temperature, washed with water, and dried. Acrylamide hydrogels were prepared with the ratio of bis-acrylamide to acrylamide to provide the target stiffness (Tse et al., 2010). Briefly, hydrogels with stiffness values of 11 or 35 kPa were prepared using 10%:0.1% and 10%:0.3% acrylamide:bis-acrylamide, respectively. Thereafter, 1 ml of gels were polymerized between the silane-coated and glutaraldehyde-treated coverslips by adding 0.5 µl of tetraethylethylenediamine (TEMED, Wako) and 5 µl of 10% ammonium persulfate (APS, Wako) until gels were formed at room temperature. The gels were then incubated overnight at 4 °C in water. Silane-coated coverslips were removed, and gels were added to 1 mM of sulfo-SANPAH (Pierce) and exposed to UV light (3000 mJ/cm^2^) using a Spectrolinker XL-1500 (Spectronics Corporation). After washing with PBS, 20 µg/ml of fibronectin (Corning) in PBS was added and incubated overnight at 4℃. The gels were washed with PBS and soaked in growth medium for at least 30 min before the cells were seeded. The Young’s moduli (*E*) of the hydrogels were measured by fitting the Hertz model using the microscopic indentation method (Lee *et al*., 2015). To visualize the surface of hydrogels, 10 μg/ml of 50 nm diameter fluorescent beads, Micromar^®^-redF (micromod), containing gel solution was polymerized between coverslips (see above). The coverslip with the hydrogel was placed on a glass-bottom dish filled with water, and a 1 mm diameter tungsten carbide sphere (AS ONE, Japan) was placed on the gel. The depth of hydrogels depression by the spheres in the z axis were measured by Z-stack confocal images (z-step size was 1 μm) using LSM710 laser-scanning confocal microscopies (Carl Zeiss) equipped with a PL Apo 10× objective lens (0.45 NA) and DPSS 561 laser diode. Each value was substituted into the formula of the Hertz model (below) to derive the *E* of the hydrogel.

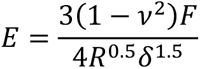

where E: elasticity of the gel; ν: Poisson’s ratio of the acrylamide gel; F; sum of the gravity and buoyancy forces of tungsten carbide sphere; R: radius of tungsten carbide sphere; δ: depth of gels depression by the spheres

### Statistical analysis

Statistical data from more than three experiments are expressed as means ± standard deviation (SD). All statistical analyses were performed using GraphPad Prism 9 (GraphPad Software, La Jolla, CA). Comparison between two groups was performed using unpaired two-tailed *t*-test, whereas comparison between more than two group was performed using one-way ANOVA, followed by Dunnett’s test to compare each treatment group with the control group, or Tukey’s test for comparisons among the treatment groups. Two-way analysis of variance (ANOVA) was used with Sidak’s test for two different conditions. Statistical significance was set at *p* < 0.05.

## Abbreviations used in this paper

ABD: actin binding domain
BioID: proximity-dependent biotin identification
BPGAP: BCH domain containing, proline-rich and Cdc42GAP-like protein
DH: Dbl homology
FAK: focal adhesion kinase
RhoGAP: Rho GTPase activating protein
RhoGEF: Rho-guanine nucleotide exchange factor
IF: intermediate filament
IP: immunoprecipitation
K8/K18: keratin 8/keratin 18
LARG: leukemia-associated RhoGEF
PA: polyacrylamide
PH: pleckstrin homology
PLEKHG4B: pleckstrin homology and RhoGEF domain containing 4B
PRG: PDZ-RhoGEF
RBD: Rho-binding domain
RGS: regulator of G protein signaling domain
SPEC: spectrin repeats

## ACKNOWLEDGMENTS

pGEX6P1-GFP-Nanobody was a gift from Kazuhisa Nakayama (Addgene plasmid # 61838 ; http://n2t.net/addgene:61838 ; RRID:Addgene_61838). Mr. Hironori Sato, Mr. Yuto Kawasaki, and Mr. Koutaro Tsuru (Tohoku university, Sendai, Japan) for technical assistance. This work was supported by Agency for Medical Research and Development (AMED) Grant Number 18gm5810015h0003 to K.O., URL: https://www.amed.go.jp/, Advanced Graduate Program for Future Medicine and Health Care, Tohoku University and JST SPRING, Grant Number JPMJSP2114 to A.K., JSPS KAKENHI Grant Numbers JP20H03248 and JP22H05618 to K.O., 21H02471 to K.M., URL: http://www.jsps.go.jp/j-grantsinaid/, and grants from the Japan Foundation for Applied Enzymology and the Uehara Memorial Foundation to K.O. A part of this study was supported by Research equipment sharing system, Tohoku University (041・TOF/TOF 5800 system; 090・LSM710 laser-scanning confocal microscopies). This work was the result of using research equipment shared in MEXT Project for promoting public utilization of advanced research infrastructure (Program for supporting construction of core facilities) Grant Number JPMXS0440600022.

